# The predictability of a lake phytoplankton community, from hours to years

**DOI:** 10.1101/230722

**Authors:** Mridul K. Thomas, Simone Fontana, Marta Reyes, Michael Kehoe, Francesco Pomati

**Author notes:** Corresponding author. Address: Centre for Ocean Life, DTU Aqua, Technical University of Denmark, Kemitorvet, 2800 Kongens Lyngby, Denmark. **Author email IDs:** Mridul K. Thomas, Simone Fontana, Marta Reyes, Michael Kehoe, Francesco Pomati.

## Abstract

Forecasting anthropogenic changes to ecological communities is one of the central challenges in ecology. However, nonlinear dependencies, biotic interactions and data limitations have limited our ability to assess how predictable communities are. Here we used a machine learning approach and environmental monitoring data (biological, physical and chemical) to assess the predictability of phytoplankton cell density in one lake across an unprecedented range of time scales. Communities were highly predictable over hours to months: model R^2^ decreased from 0. 89 at 4 hours to 0.75 at 1 month, and in a long-term dataset lacking fine spatial resolution, from 0.46 at 1 month to 0.32 at 10 years. When cyanobacterial and eukaryotic algal cell density were examined separately, model-inferred environmental growth dependencies matched laboratory studies, and suggested novel trade-offs governing their competition. High-frequency monitoring and machine learning can help elucidate the mechanisms underlying ecological dynamics and set prediction targets for process-based models.

## Introduction

Forecasting how environmental change will alter communities and ecosystems is perhaps the most important task facing ecologists today, and a tremendous challenge to our ecological understanding (Mouquet et al. 2015, Petchey et al. 2015, Houlahan et al. 2016). Nonlinear relationships (such as between temperature and most biological processes), stochasticity and sensitive dependence on initial conditions are sources of uncertainty that community ecology shares with other predictive disciplines, such as climate science. However, ecology additionally has to grapple with biotic interactions in complex food webs, evolutionary change, and a paucity of data with which to assess predictive power and refine models (Magurran et al. 2010). Therefore, with few exceptions (notably in disease ecology, e.g. Axelsen et al. 2015), we do not know how predictable ecological communities are (i.e. how strong the association between present and future system states is, a proxy for forecast ability). Quantifying this would allow us to understand the time scale over which we can provide actionable input for management and legislative decision-making, recently termed the *ecological forecast horizon* (Petchey et al. 2015).

To make accurate long-term forecasts, ecology needs to develop process-based forecasts akin to those prevalent in climate science. Correlational approaches based on present conditions and abundances are likely to make inaccurate forecasts over decadal time-scales because patterns of environmental covariation will change in the future (Williams et al. 2007). Process-based models avoid this problem but are a challenge to design because of their complexity. This arises from of a lack of knowledge of the functional forms (or shape) relating population/community change to environmental factors, and a lack of data with which to parameterise them (Kremer et al. 2016). The scale of this challenge is highlighted by recent work showing that complex, highly nonlinear interactions between abiotic factors are a regular feature of physiological and ecological processes (Zhu et al. 2016, Zimmer et al. 2016, Edwards et al. 2016, Thomas et al. 2017). Designing process-based models using a traditional approach may require extensive experimental work examining high-dimensional interactions. High-throughput screening technologies are helping to address this problem. But in many cases, a traditional experimental approach to understanding interactions (i.e. through multidimensional factorial experiments) may not be realistic given present funding and experimental constraints.

Machine learning (ML) algorithms offer us an alternative path towards the creation of these process-based models. When presented with complex environmental datasets, ML allows us to avoid the most important constraints inherent in traditional statistical approaches (*a priori* specification of functional forms, interactions and error distributions). Despite relying on underlying correlations, ML algorithms can improve substantially on traditional correlative analyses (Rivero-Calle et al. 2015, Kehoe et al. 2012, 2016). They can be used on complex datasets to assess associations (Rivero-Calle et al. 2015) and quantify predictability in the absence of the knowledge needed for a process-based model (Ewers et al. 2017). Even more importantly, they can be used to *infer* the functional forms and interactions needed to develop process-based models. This approach will require large datasets, but as ecology enters the ‘big data’ era, acquiring this is becoming feasible for many systems. As the cost of data acquisition continues to decrease, ML may prove a more efficient approach (relative to high-dimensional factorial experiments) to assessing community predictability and understanding the drivers of complex ecological dynamics.

Natural communities of microbes such as phytoplankton are vital parts of most biogeochemical cycles and food webs (Field et al. 1998, Falkowski et al. 1998), and so assessing their predictability is especially important. Phytoplankton have generation times on the order of a day, and respond extremely rapidly to environmental change: shifts on time-scales of minutes to hours are sufficient to elicit physiological and ecological changes (Goldman & Glibert 1982, Demers et al. 1991, Hemme et al. 2014). Despite this sensitivity to environmental conditions, we do not know the time-scales over which phytoplankton community dynamics may be predicted.

Historically, most plankton monitoring campaigns have measured the community at coarse time-scales of once to twice a month (Jochimsen et al. 2012), amounting to tens of generations. These efforts have helped us understand broad changes driven by eutrophication and environmental warming (Pomati et al. 2012, Jochimsen et al. 2012), helping to make the case for policies limiting further changes. However, with rare exceptions (notably Hunter-Cevera et al. 2014, 2016), plankton monitoring efforts have not captured data needed to accurately assess community predictability across time scales. High-frequency monitoring campaigns that sample communities and environmental drivers on sub-daily time-scales can partly address this (Pomati et al. 2011, Merel 2013, Pomati et al. 2013, Hunter-Cevera et al. 2014, 2016), filling in pieces of the picture that coarser long-term datasets have hinted at. They also provide us with the quantity of data needed to profitably employ ML tools.

We quantified the predictability of phytoplankton cell density over time scales ranging from 4 hours to 10 years, or approximately 10^−1^ to 10^3^ generations. Cell density, or the abundance of phytoplankton cells per unit volume, is the most important parameter characterising the phytoplankton community. It is a strong proxy for phytoplankton biomass (including in our system, Fig. S1) and for primary productivity, an important ecosystem property. We characterised predictability of cell density in Greifensee, a meso-eutrophic peri-alpine lake in Switzerland. The Greifensee plankton community, chemistry and physics have been monitored for >30 years, during which it has seen dramatic changes in biology as a result of eutrophication and re-oligotrophication (Bürgi et al. 2003). We make use of two complementary datasets examining the Greifensee phytoplankton community: (1) High-frequency data from monitoring campaigns carried out in the summer and autumn of 2014 and 2015. Cell density was measured at 6 different depths (Fig. 1) using scanning flow cytometry (SFCM), and environmental data were also collected (Table S1). (2) A long-term time series created from monthly measurements of depth-integrated phytoplankton measurements (Fig. 1), as well as associated environmental factors (Table S2). Although the datasets differ in methodology and size, they measure substantially similar biological, physical and chemical factors (Tables S1, S2). Given the length of the two datasets and their sampling frequency and location, we are able to directly compare predictability in the two datasets at a time lag of 1 month.

**Fig. 1.**
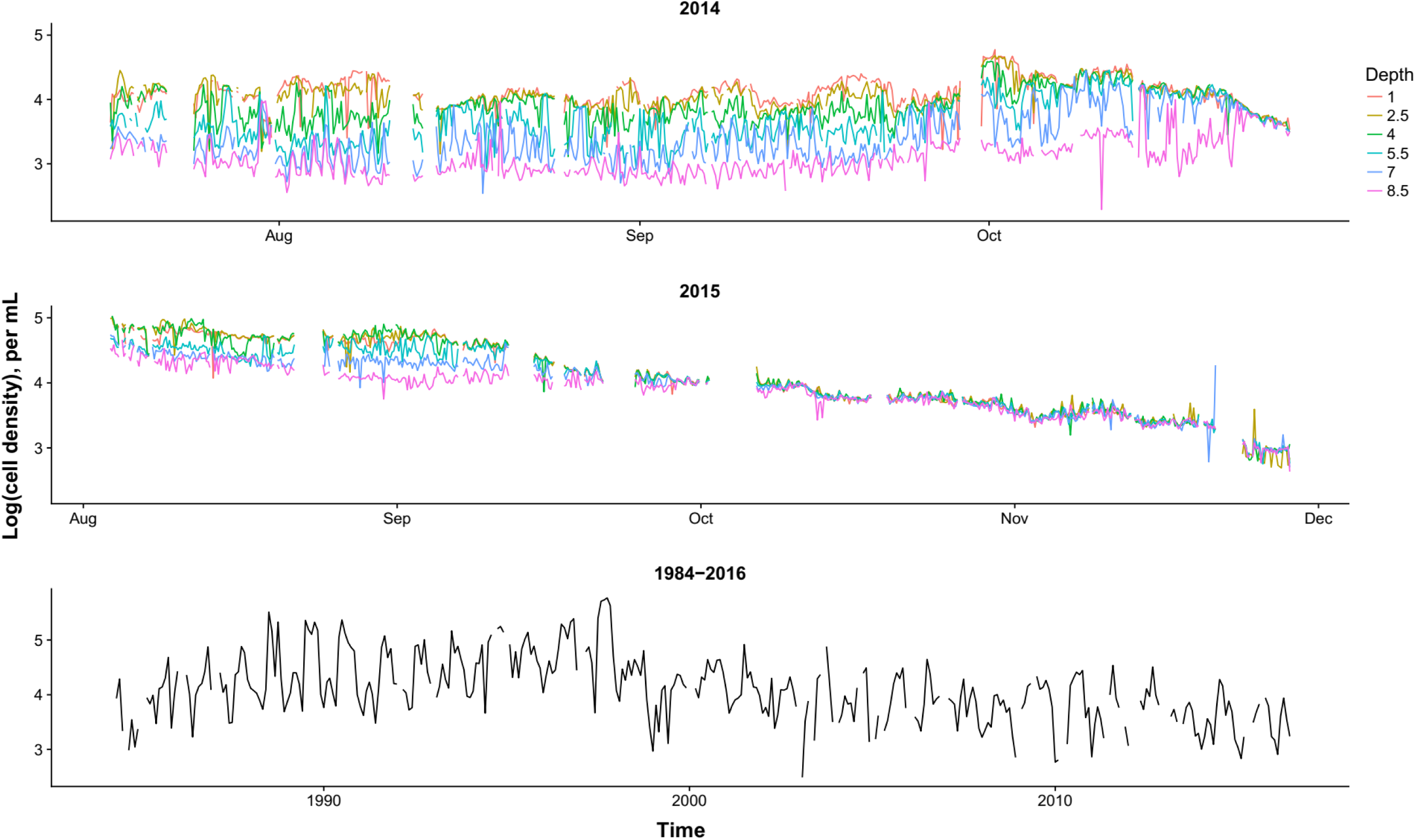
Dynamics of cell density of the total phytoplankton community, in both the high-frequency and long-term datasets from Greifensee. High-frequency measurements were made every 4 hours in summer-fall 2014 and 2015, at six depths. Long-term measurements were made monthly from 1984 to 2016 and were integrated over the top 20m. Note that X-axes are on different scales in each panel. Y-axes are identical for the top two panels but differ for the third.

In addition to community density, ecology aims to predict the dynamics of functional groups and taxa. Especially because toxic cyanobacterial blooms are a major health concern (Chorus & Bartram 1999, Paerl & Huisman 2009, Paerl et al. 2011, Merel et al. 2013), we also assessed the predictability of cyanobacterial cell density over these time scales, and the drivers of competitive dynamics between cyanobacteria and eukaryotic phytoplankton (Fig. 2). Eukaryotic densities have remained relatively stable in Greifensee since the 1980s, while average cyanobacterial densities first increased 100-fold during eutrophication and then decreased by a similar amount over this time period as a result of re-oligotrophication (Fig. 2). Understanding the drivers of growth and competition between these two broad phytoplankton groups can help us refine process-based models of water quality, with important implications for the management of aquatic ecosystem services.

**Fig. 2.**
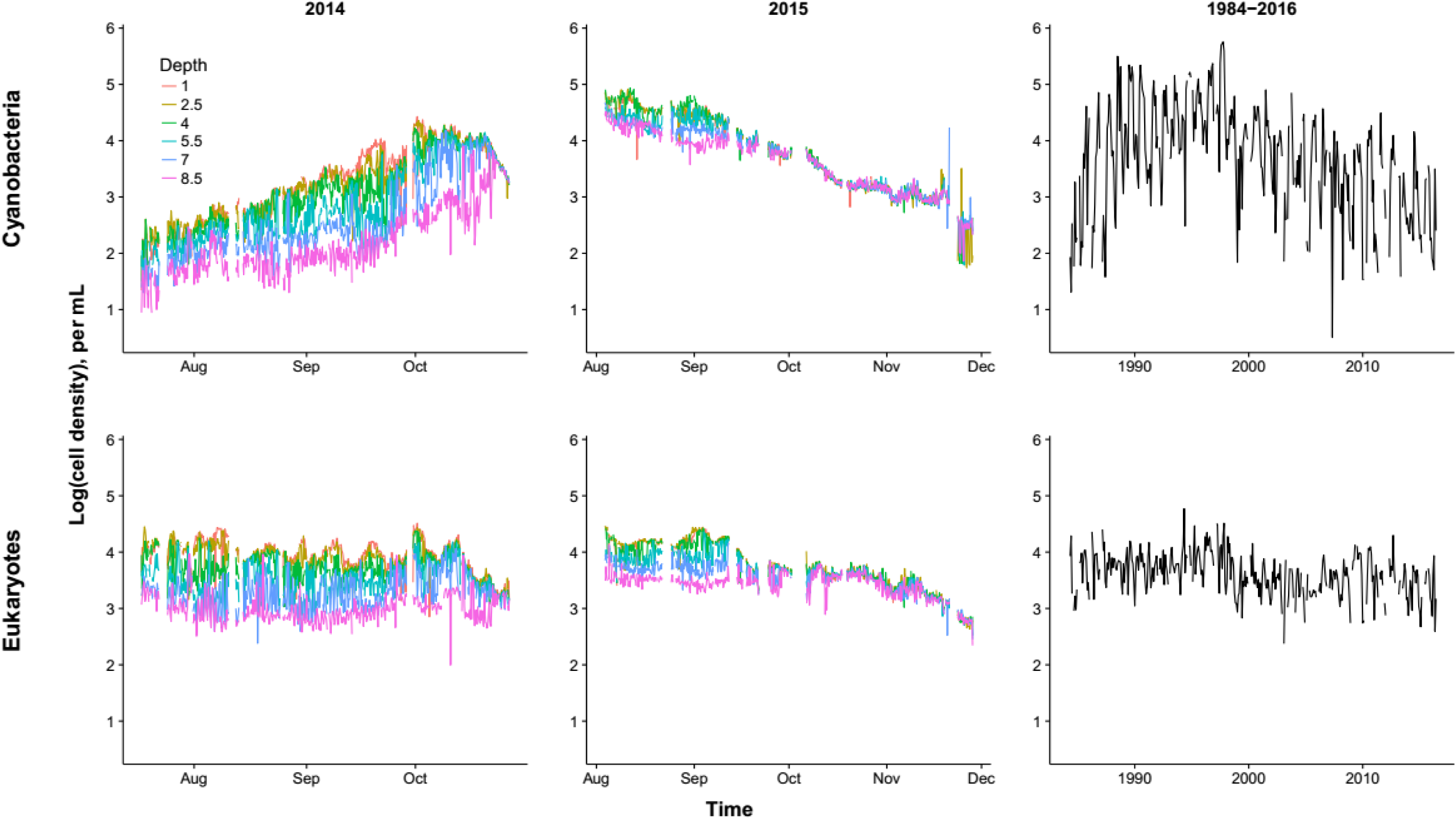
Dynamics of cell density of the cyanobacteria and eukaryotic phytoplankton, in both the high-frequency and long-term datasets from Greifensee. High-frequency measurements were made every 4 hours in summer-fall 2014 and 2015, at six depths. Long-term measurements were made monthly from 1984 to 2016 and were integrated over the top 20m. Note that X-and Y-axes are on different scales in each column.

## Methods

### I. Overview

Greifensee is a peri-alpine lake in Switzerland (47.35°N, 8.68°E) with a documented history of eutrophication and re-oligotrophication (Bürgi et al. 2003). The lake is currently meso-eutrophic, 32 m deep at its deepest point and just over 20 m deep at the sampling locations used for both datasets.

We use two datasets in this study: a high-frequency dataset consisting of measurements every 4 hours during the summer and autumn of 2014 and 2015 using the automated monitoring station Aquaprobe (Pomati et al. 2011), and a long-term dataset consisting of monthly measurements from March 1984 to June 2016. In both cases, important environmental data (both abiotic and biotic) was collected simultaneously near the middle of the lake, allowing similar analyses to be conducted and thereby enabling comparisons. However, the datasets differ in important ways:

i. The high-frequency dataset involved measurements by SFCM, while the long-term dataset involved microscopy measurements. Therefore, sampling effort and density assessment methods differ.
ii. The high-frequency dataset consists of measurements at six specific depths (1.0, 2.5, 4.0, 5.5, 7.0 and 8.5m). Abiotic environmental data were also estimated at the same depth as the collected sample. In contrast, the long-term dataset consists of integrated phytoplankton measurements across the top 20m of the lake. Abiotic measurements were not integrated, but collected at specific depths (except for light, which is a surface estimate), and so we calculated the maximum and minimum value of each abiotic factor in the top 20m for use as our predictors.
iii. The high-frequency dataset consists of 7161 measurements, while the long-term dataset contains 383 measurements.

### II. High-frequency dataset generation

#### 1. Scanning flow cytometry description

We used a scanning flow cytometer, the CytoSense (http://www.cytobuoy.com), to quantify the density of the total phytoplankton community as well as its cyanobacterial and eukaryotic algal fractions (estimated densities are strongly correlated with estimates from microscopy, Fig. S2). The CytoSense characterizes the scattering and pigment fluorescence of individual phytoplankton cells. It measures cells and colonies across a large proportion of the phytoplankton length range, between approximately 2 μm and 1 mm in length. Particles that enter the system cross two coherent 15mW solid-state lasers. The instrument’s laser and sensor wavelengths are designed to target the fluorescence signals primarily from chlorophyll-a and phycocyanin, but also capture signals from phycoerythrin and carotenoids. We used two different instruments 2014 and 2015, with small differences in configuration. Instrument settings and data processing steps may be found in the supplementary information.

#### 2. SFCM field sampling procedure

All samples were collected from a floating platform Aquaprobe (Pomati et al. 2011) near the middle of the lake (47.3663°N, 8.665°E). Every four hours, water was sampled automatically at each of the 6 depths (as described in Pomati et al. 2011). Water samples were pumped into a 150 mL sampling chamber at the surface through a tube with a 0.6-cm diameter opening. The sampling chamber was flushed with water from the sampling depth three to five times over 2 minutes before the CytoSense collected a subsample of up to 500 μL for measurement.

#### 3. Environmental factors

The full list of environmental parameters is found in Table S1. We measured temperature, conductivity and irradiance at all depths. We also collected weekly depth-specific samples for dissolved nutrient (nitrate, phosphate, ammonium) concentration estimation, and integrated measurements of size-fractionated zooplankton. We also monitored meteorological factors including wind speed and rainfall, and made use of additional data provided by the Office of Waste, Water, Energy and Air (AWEL) of Canton Zürich on water inflow (including flow rate, temperature and nutrient concentrations) into the lake. Detail of sampling procedures, instruments used and measurement methodology may be found in the supplementary information.

### III. Long-term dataset

#### 1. Field sampling procedure

Approximately every month, water samples were collected for physical, chemical and biological measurements near the centre of the lake (47.3525°N, 8.6748°E), approximately 1.5 km from the floating platform used for the high-frequency measurements. For microscopic counts of the phytoplankton community, an integrated water sample was collected over the upper 20 m of the water column with a Schröder sampler (Bürgi et al. 2003).

#### 2. Environmental factors

The full list of parameters is found in Table S2. To measure chemical and physical parameters, water samples were collected every 2.5 m over the whole water column, at the same location and on the same dates, and were analysed using standard limnological methods (Rice et al. 2012). Integrated zooplankton samples were collected over the upper 20 m of the water column. We also made use of a publicly available surface irradiance dataset (Schulz et al. 2008, Müller et al. 2015) to estimate the monthly-averaged irradiance at the water surface based on interpolated estimates from a location approximately 2 km from the sampling location (47.35°N 8.65°E). More details about sampling procedures, instruments used and measurement methodology may be found in Bürgi et al. (2003).

#### 3. Data processing

For every time point, we calculated the maximum and minimum value of every depth-specific parameter (such as phosphate concentration) across the entire water column, and used these for subsequent analyses. Additionally, samples were not collected on the same day every month and, in rare cases, more than one sample was collected in a month. We therefore aggregated measurements by rounding to the nearest month and then averaged duplicate values.

### IV. Machine learning analyses

#### 1. Random forests overview

Random forests (RFs) are a robust ML tool comprising ensembles of regression trees (or classification trees) (Breiman 1999). In each regression ‘tree’ within the random ‘forest’, a randomly selected subset of the data is recursively partitioned based on the most strongly associated predictor. At each node, a random subset of the total number of predictors is considered for partitioning. The final tree ‘prediction’ for new data is given by the average value of the data within each branch of the tree. By aggregating predications across trees, RFs are able to reproduce arbitrarily complex shapes patterns without *a priori* functional form specification.

We took advantage of three features that make RFs a flexible and useful tool for examining ecological systems: *permutation importance*, easy quantification of *partial effects* of individual predictors, and *out-of-bag prediction*.

i. The importance of each predictor in a RF is assessed by permuting the predictor across all trees in the forest and quantifying the resulting change in the forest’s error rate. More important predictors lead to a greater increase in error when permuted.
ii. The partial effect of any single predictor on the dependent variable can also be quantified, allowing us to examine the functional form of the relationship (which may be arbitrarily nonlinear, though non-bifurcating).
iii. Out-of-bag (OOB) prediction allows us to make accurate estimates of error rate and goodness of fit (model R^2^) via a process akin to cross validation (Breiman 1999). Each data point is present in the training data of only a subset of all ‘trees’ that comprise the ‘forest’. Therefore, the value of every point may be predicted using the trees that have not been trained with it. The ‘OOB prediction error’, or mean difference between the OOB predictions and the true value of all points in the dataset (see Fig. S3, S4 for examples using our data) can be used to quantify the RF’s predictive ability through a pseudo-R^2^:

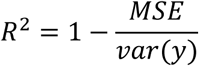

where *MSE* is the mean squared error of the OOB predictions when compared to the true values, and *var(y)* is the variance in the dependent variable. As in the case of a standard R^2^, a pseudo-R^2^ has an upper bound of 1, indicating perfect model performance. However, note that unlike a standard R^2^, there is no lower bound. It is possible for the pseudo-R^2^ to be negative, if *MSE* > *var(y)*. This may be interpreted as saying that the model prediction is worse than the mean value of the dependent variable in the entire dataset. In our analyses, we saw low, negative values of pseudo-R^2^ in a small number of cases; we rounded these values to zero to avoid confusion, while noting this in the figure captions.

#### 2. Data pre-processing

In our analyses, we omitted: i) all entries cases where the dependent variable was missing, and ii) the predictors *sampling depth* and *sampling time*. We omitted the latter in order to accurately estimate the effects of predictors that covary with depth and time on cell density. In other words, we believe that gradients in light, temperature and nutrients should characterise most of the relevant information contained within depth and time.

#### 3. Quantifying predictability

We quantified the *predictability* of log cell density of the total phytoplankton community, and the cyanobacterial and eukaryotic fractions, using the pseudo-R^2^ calculated based on the OOB predictions of the fitted model (see details above). We estimated predictability at time lags ranging from 4 hours to 1 month in the high-frequency dataset, and from 1 month to 10 years in the long-term dataset. For every time lag, we fit two models, predicting log cell density using: 1) only log cell density at the specified time lag, and 2) both log cell density and environmental parameters at the specified time lag. E.g. our simplest model considering a time lag of four hours predicted log cell density at all time points using only log cell density from the measurement four hours previously.

#### 4. Predictor importance

We assessed the importance of predictors at all time lags using the change in model error rate when the predictor values were permuted.

#### 5. Partial effects of environment on growth

We quantified the model-inferred effects of environmental factors on cyanobacteria and eukaryotes. Instead of examining the effects of these predictors on log density, we instead examined how they influence the population growth rate (i.e. specific growth rate, day^−1^, the rate of change in density between successive time points). We did this to facilitate comparison between the partial effects in our field dataset and extensive prior laboratory findings for the same predictors. However, the functional forms remained highly similar to the model explaining log density.

We focussed on two factors that are known to influence phytoplankton growth (Litchman & Klausmeier 2008) and were identified as important in our analyses: light and temperature.

Because laboratory studies typically measure the effects of environmental factors on population growth rate *per day*, we multiplied the estimates of growth rate over four hours by 6 to express them in the same units. We then fitted RFs to these population growth rates using environmental parameters at a 4-hour lag and estimated their partial effects. Note that we omitted log density as a predictor, but model structure was otherwise identical to those previously described.

Though we were also interested in the effects of dissolved nitrate, phosphate and N:P ratio, we had less well-resolved data for these predictors that limited the power of analyses relating to these factors.

#### 5. Model fitting and settings

All analyses were done in the *R* statistical environment v3.3.3 (R Core Team 2017) using the package *randomforestSRC* (Ishwaran & Kogalur 2007, Ishwaran & Kogalur 2017). We used 2000 trees for every forest, and set the number of predictors to be considered at each node to be one-third of the total number of predictors. Missing data among the predictors were imputed for the purpose of model fitting, but imputed values were not used for predictor importance assessment (Ishwaran & Kogalur 2007, Ishwaran & Kogalur 2017).

## Results

Phytoplankton cell density was highly predictable on time scales of hours to months. In our high-frequency dataset, pseudo-R^2^ of the RF models trained with cell density and environmental data decreased from 0.89 at a 4 hour lag to 0.74 at a lag of 1 month (Fig. 3). The model using only cell density as a predictor had a lower R^2^ at all time lags. As time lag increased, including environmental data led to larger improvements in predictability: the difference in R^2^ between the two models was 0.03 at a 4 hour lag, and ten times higher (0.30) at a lag of 1 month (Fig. 3). In the long-term dataset, R^2^ of the model trained with cell density and environmental data decreased from 0.46 at a time lag of 1 month to 0.35 at 6 months, after which it remained relatively stable (Fig. 3). R^2^ in the density-only model was lower at all time lags.

**Fig. 3.**
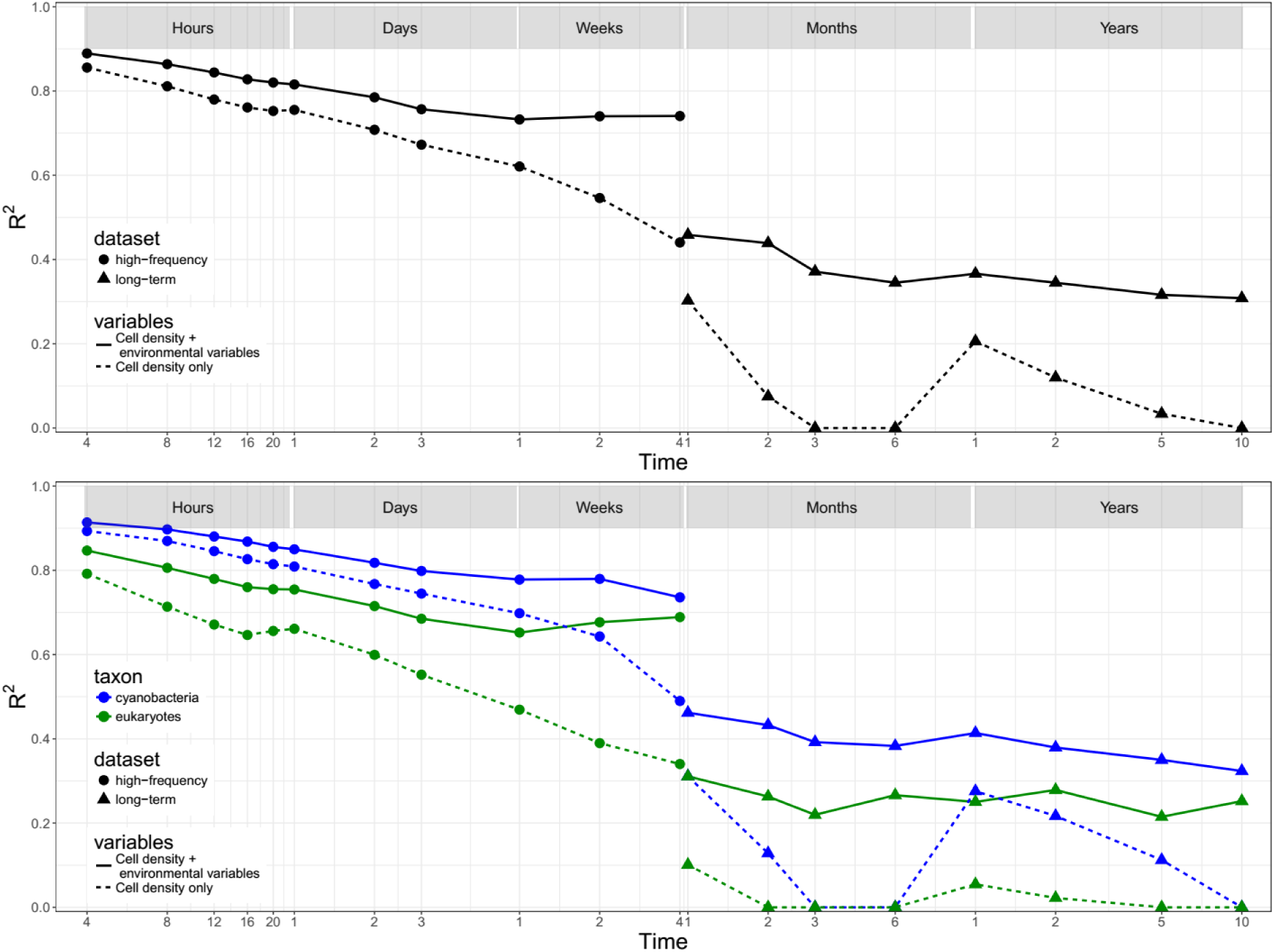
Decline in predictability of the phytoplankton community with time, characterised by the random forest pseudo-R^2^. The predictive contribution of environmental information increased with increasing time lag (distance between solid and dashed lines increases). Cyanobacteria were consistently more predictable than eukaryotes. Despite overlap between high-frequency and long-term datasets at a time lag of 1 month, there is a decline in predictability likely driven by the lack of depth resolution in plankton and environmental data in the long-term dataset. The spike in R^2^ of the ‘cell density only’ models at 1 year reflects strong annual cycles in density. Note that pseudo-R^2^ values can go negative (see Methods), and we rounded a few slightly negative values up to zero. We present the same results in terms of change in Mean Absolute Error with increasing time lag in Fig. S5.

Aside from cell density, which was the strongest predictor at all time lags in our high-frequency dataset, the most important predictors were light, temperature and thermocline depth, itself an indirect effect of temperature (Fig. 4). Light and temperature were also most important in our long-term dataset on time scales of months (Fig. 4). At time-scales of years, dissolved phosphorus and zooplankton density become more important.

**Fig. 4.**
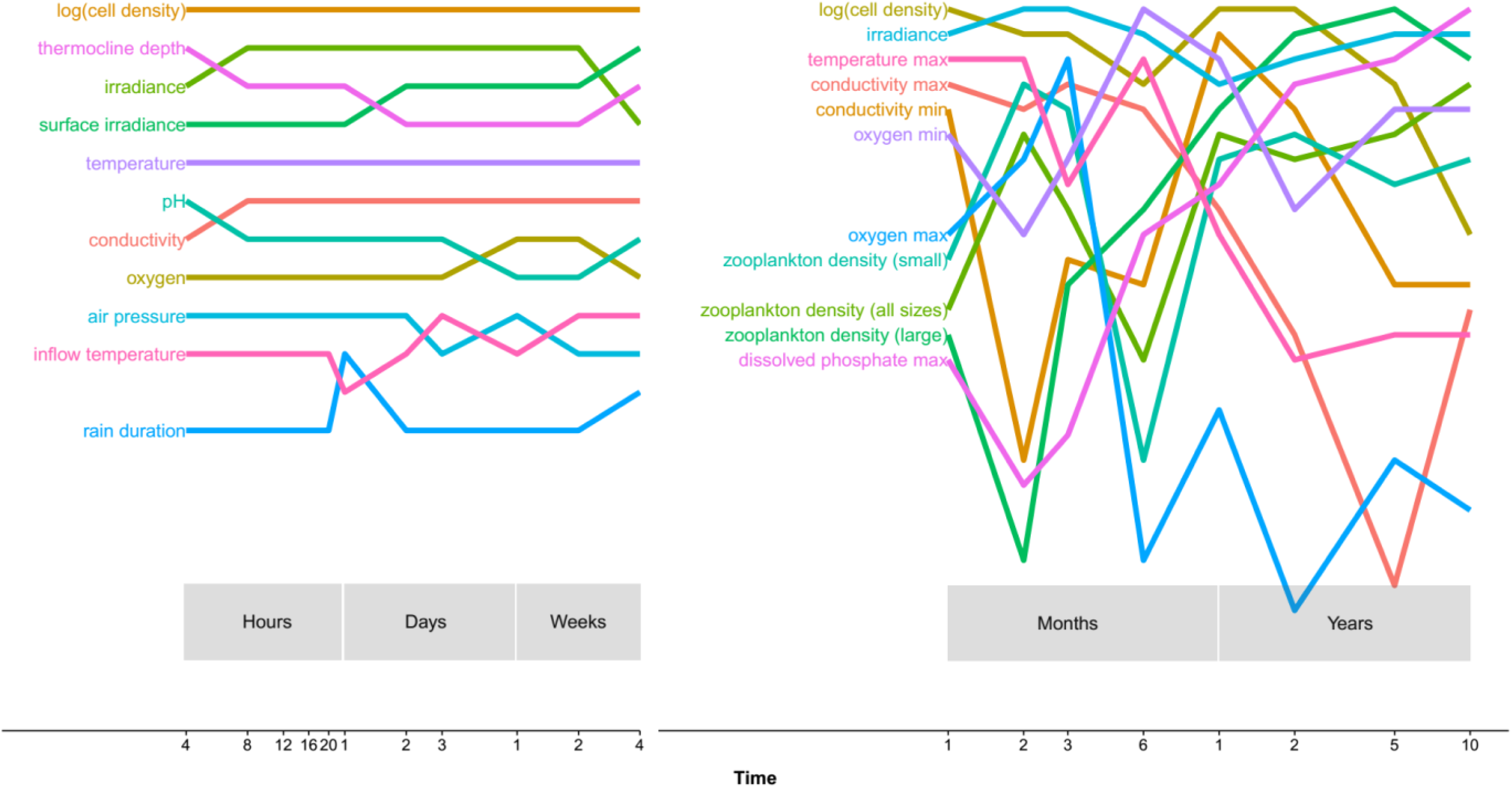
The most important predictors of phytoplankton cell density at different time lags, ordered by descending rank. Light and temperature (directly, or indirectly through thermocline depth) were important predictors at most time scales. In the long-term dataset, phosphorus and zooplankton density become highly important predictors at time scales of >1 year. Only the most important variables are shown here, for legibility (the top 5 predictors contribute >80% of the predictive power in most cases). In the high-frequency dataset, only variables that are in the top 10 most important for at least one time lag are shown, while in the long-term dataset we show only variables that appear in the top 5 most important at least once. See Tables S4 and S5 for the importance of all variables tested.

Cyanobacteria were more predictable than eukaryotes at all time scales, in both high-frequency and long-term datasets (Fig. 3). Model R^2^ for cyanobacteria was consistently higher that than for eukaryotes by approximately 5-20 percentage points (Fig. 3), in both types of models (cell density only and cell density with environmental data).

To motivate the development of process-based models of phytoplankton competition, we also examined the partial effects of environmental factors on the growth rate of cyanobacteria and eukaryotic algae (Fig. 5). Temperature and light, the strongest predictors in our dataset, showed biologically realistic nonlinear patterns. Additionally, in both cases, each group dominated a region of parameter space, suggesting the presence of trade-offs in performance.

**Fig. 5.**
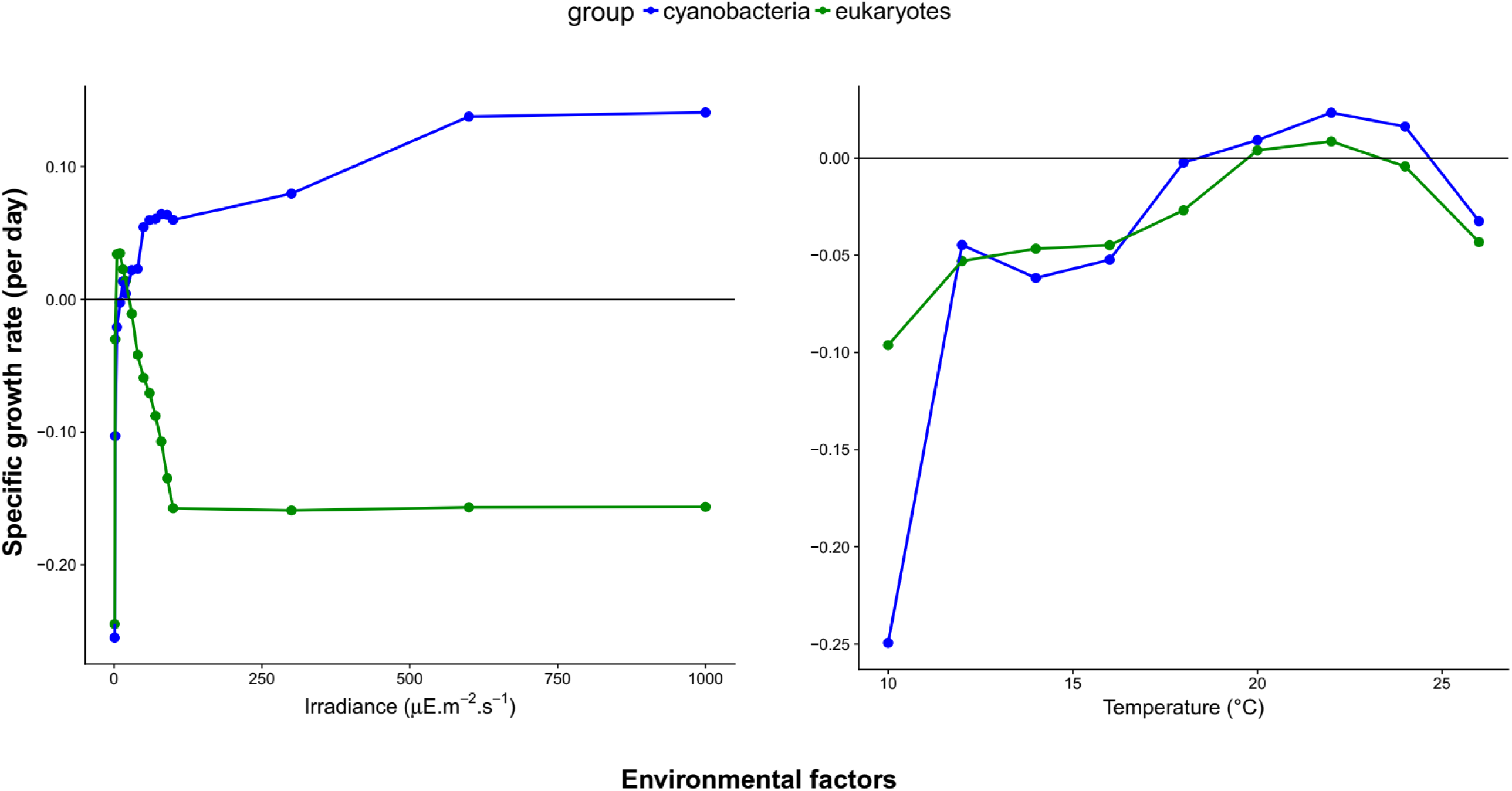
Partial effects of important environmental variables on the population growth rates of cyanobacteria and eukaryotes, based on an RF model with a 4-hour time lag. The patterns reveal environmental dependencies consistent with lab experiments and suggest trade-offs with important ecological implications. Cyanobacteria appear to benefit from high light and high temperature. Wholly negative partial effects reflect the fact that in 2015, the community was decreasing through the majority of the monitoring campaign. Therefore, differences in shape and magnitude are highly informative, but the absolute estimates are only indicative. This is especially true because interactions between variables are captured by the complete forest prediction, but are not visible in single-dimension partial effects plots. Note that Y-axes are on different scales.

## Discussion

Assessing the predictability of natural communities is crucial if we are to develop forecasts of how ecosystems will be altered by anthropogenic environmental change (Petchey et al. 2015, Mouquet et al. 2015, Houlahan et al. 2016). However, our ability to predict community dynamics has been limited by our understanding of environmental dependencies and biotic interactions (McGill et al. 2006). Our results suggest that lake phytoplankton communities are highly predictable over time scales of hours to months, approximately 10^−1^ to 10^2^ generations, and possibly longer (Figs. 3, S5). Our approach quantifies the decline in predictability with increasing time lag, identifies the predictors that contribute to predictive power, and points towards realistic trade-offs and parameterisations through the examination of partial effects. Together, these can inform the development of process-based models, set targets for their forecasts to achieve, and identify a forecast horizon for adaptive management strategies. This is especially true in the case of cyanobacteria, which are a threat to human health and aquatic ecosystem services because of toxin production, and are believed to be hard to forecast (Chorus & Bartram 1999, Paerl & Huisman 2009, Paerl et al. 2011, Merel et al. 2013). We find cyanobacterial densities to be consistently more predictable than those of eukaryotes (Fig. 3).

As our understanding of ecological processes improves, the limits to predictability of ecological systems will be determined by more fundamental constraints such as stochasticity, and sensitive dependence on initial conditions. Despite these forces, we find strong, ecologically important environmental forcing in a natural system across a range of time scales (Figs. 3-5). Consequently, we believe that process-based models are very likely to provide us with useful predictions over medium-to-long time scales. In other words, we believe that despite the complexity of phytoplankton communities, the ecological forecast horizon (Petchey et al. 2015) is sufficiently distant for ecologists to provide useful input into adaptive management strategies. Note that we do not quantify a specific horizon here because this requires the specification of a (arbitrary) forecast threshold; if desired, readers may choose these thresholds for themselves and identify the resulting forecast horizon from Fig. 3.

Light and temperature were strongly predictive of phytoplankton dynamics across time scales (Fig. 4), consistent with existing ecological understanding (Litchman & Klausmeier 2008). We also found that zooplankton density and dissolved phosphorus concentrations become highly predictive on time scales longer than a year, consistent with an ongoing, multidecadal decrease in phosphorus and biomass in Greifensee (Buergi et al. 2003). This identification of variables that are known to play a major role in phytoplankton ecology strengthens our confidence in the relationships underlying our metric of predictability. However, we note that predictor importance – while a useful tool – is sensitive to missing data patterns. Our estimates therefore understate the importance of two major groups of predictors in our high frequency data: nutrients and zooplankton density. Unlike most other predictors that were measured every four hours, these were measured weekly in 2014 and twice a week in 2015 (Table S1). To partially correct for this difference, we also assessed the relative importance of all predictors when these were interpolated using generalised additive models (GAMs) (Fig. S6). Models with interpolated nutrients and zooplankton predictors had marginally higher R^2^ values and these predictors rose considerably in importance, especially dissolved nitrogen. We believe that these results are noteworthy, but choose not to focus on them here because we are unable to validate the interpolated estimates.

Importantly, the predictive power of environmental factors in our models arises from nonlinear dependencies that are consistent with causal relationships established through lab studies (Fig. 5; Litchman & Klausmeier 2008). Light, one of the most important predictors, has a partial effect on growth that is a saturating function for cyanobacteria and a right-skewed unimodal function for eukaryotes (Fig. 5); these are the only shapes consistent with laboratory measurements of light-dependent growth (Eilers & Peeters 1988, Edwards et al. 2015). The partial effect of temperature is an increasing function and possibly a left-skewed unimodal curve, consistent with prior eco-physiological findings, including in phytoplankton (Kingsolver 2009, Thomas et al. 2012, Thomas et al. 2016). This concordance between controlled lab studies and ML-derived field patterns increases our confidence in the suitability of this ML approach, and suggests that the relationships we have uncovered are likely to be useful in guiding process-based model creation. Furthermore, it suggests that ML approaches may be used to discover novel ecological patterns. This is particularly important in the case of interactions between factors, which presently require labour-intensive and expensive multifactorial experiments to understand.

The partial effects that we show here (Fig. 5) point towards trade-offs that could enable the coexistence of cyanobacteria and eukaryotes. Cyanobacteria appear to benefit from high light intensity and high temperature, while eukaryotes have a growth advantage in the converse conditions. Therefore, temporal heterogeneity in one or both of these dimensions could allow for the maintenance of both these groups (Chesson 2000). Cyanobacteria do possess higher optimal temperatures for growth than eukaryotic phytoplankton at temperate latitudes (Thomas et al. 2016), consistent with the temperature-dependence we see (Fig. 5). The apparent trade-off between growth at high and low light intensities was not seen in a synthesis of lab-measured light traits (Schwaderer et al. 2011), but at present, measurements are available only from a small number of species and may be influenced by interactions with other factors. Laboratory data on a broader range of species and under a greater range of conditions will be needed to resolve this discrepancy. If true, the pattern we observe in the field also suggests an explanation underlying the formation of surface scums by cyanobacteria through buoyancy regulation (Paerl et al. 2011, Carey et al. 2012). Scum formation - important due to the negative impact on lake ecosystem services - is consistent with a cyanobacterial benefit from higher irradiance. In contrast, eukaryotes appear to have a lower optimal irradiance and might experience photo degradation from surface growth. These observations offer an example of the insights that may be gained through a combination of high-frequency monitoring and machine learning.

Our models may understate the long-term predictability of the phytoplankton community. The difference in predictability between high-frequency and long-term datasets at a time lag of 1 month suggests that if a similar methodology was followed in the long-term dataset, reasonably high R^2^ values may have been obtained over time scales of years, not just months. This difference is driven by several factors: 1) our high-frequency dataset includes measurements of both phytoplankton and environmental factors at specific depths, as opposed to integrated values across the water column as in the long-term dataset, 2) the high-frequency dataset has more than an order of magnitude more data points (7161 vs. 383) with which to train the machine learning algorithm, and 3) the long-term dataset explores a far greater range of parameter space in temperature, nutrient concentration, zooplankton density and unmeasured factors. Of the three, we believe depth-specific sampling may be the major factor, as the difference in model R^2^ at a lag of one month is <15% in the case of the cell density-only models, and 30% in the models with both cell density and environmental factors (Fig. 3). However, we also note that in more complex systems where migration is a larger factor – such as coastal and open-ocean communities – predictability may be lower unless physical circulation patterns are highly predictable as well.

It is important to note that although pseudo-R^2^ provides estimates of predictability that are robust (Breiman 1999), we have not assessed a true *forecast*, in which error is allowed to compound through time. This approach can in principle be used to make a forecast, but we chose not to do so because of large changes in environmental conditions towards the end of the 2014 and 2015 monitoring seasons. Attempting to forecast would require us to predict in conditions well outside those that the model was trained on, where it will inevitably perform poorly. Despite this limitation, we believe that out-of-bag error is a useful proxy for forecast error: the realistic environmental dependencies (Fig. 5) highlight that we are uncovering the mechanisms underpinning ecological dynamics. In the future, a broader sampling of parameter space (through a year-round monitoring campaign) should allow us to make and test true forecast skill.

We have shown that high-frequency environmental monitoring and machine learning approaches can be usefully employed to uncover patterns in complex ecological communities, to assess the predictability of these communities, and to uncover dependencies that can then be incorporated into process-based models of communities and ecosystems. This can help us address fundamental questions in ecology: What are the drivers of ecological processes and how does this change through time? How large of an effect does environmental and demographic stochasticity have on communities? What are the dominant trade-offs that maintain diversity in natural systems and how do they operate in dynamic environments? But perhaps more importantly, it can allow us to improve our forecasts of ecological systems, fulfilling a fundamental obligation that ecology owes to society.

## Acknowledgements

This work was funded by Swiss National Science Foundation grants CRSII2_147654 and 31003A_144053. We thank: the Office of Waste, Water, Energy and Air (AWEL) of Canton Zürich for providing permission for *in situ* monitoring and data on water inflow into Greifensee; the lab groups of H. R. Buergi and P. Spaak for data collection and access to the long-term dataset; Esther Keller for assistance with microscopy; Dany Steiner, Hannele Penson and Christian Ebi for help with field work and equipment maintenance; Idronaut and Cytobuoy for support during monitoring campaigns; and Colin T. Kremer for helpful comments on the manuscript.

## Statement of authorship

MKT & FP conceived the study. MKT, SF, MR & FP collected the data. MKT analysed the data. MKT wrote the manuscript with substantial input from FP, SF, MK & MR.

## References

Axelsen JB, Yaari R, Grenfell BT, Stone L (2014) Multiannual forecasting of seasonal influenza dynamics reveals climatic and evolutionary drivers. PNAS, 111, 9538–9542.

Breiman L (1999) Random Forests. Machine Learning, 45, 1–35.

Bürgi HR, Bührer H, Keller B (2003) Long-Term Changes in Functional Properties and Biodiversity of Plankton in Lake Greifensee (Switzerland) in Response to Phosphorus Reduction. Aquatic Ecosystem Health & Management, 6, 147–158.

Carey CC, Ibelings BW, Hoffmann EP, Hamilton DP, Brookes JD (2012) Eco-physiological adaptations that favour freshwater cyanobacteria in a changing climate. Water Research, 46, 1394–1407.

Chorus I, Bartram J (1999) Toxic Cyanobacteria in Water: A Guide to Their Public Health Consequences, Monitoring, and Management. E & FN Spon, 416 pp.

Conley DJ, Paerl HW, Howarth RW et al. (2009) Controlling eutrophication: Nitrogen and Phosphorus. Science, 323, 1014–1015.

Demers S, Roy S, Gagnon R, Vignault C (1991) Rapid light-induced changes in cell fluorescence and in xanthophyll-cycle pigments of Alexandrium excavatum (Dinophyceae) and Thalassiosira pseudonana (Bacillario-phyceae): a photo-protection mechanism. Marine Ecology Progress Series, 76, 185–193.

Edwards KF, Thomas MK, Klausmeier CA, Litchman E (2015) Light and growth in marine phytoplankton: allometric, taxonomic, and environmental variation. Limnology and Oceanography, 60, 540–552.

Edwards KF, Thomas MK, Klausmeier CA, Litchman E (2016) Phytoplankton growth and the interaction of light and temperature: A synthesis at the species and community level. Limnology and Oceanography, 61, 1232–1244.

Eilers PHC, Peeters JCH (1988) A model for the relationship between light intensity and the rate of photosynthesis in phytoplankton. Ecological Modelling, 42, 199–215.

Ewers RM, Andrade A, Laurance SG, Camargo JL, Lovejoy TE, Laurance WF (2017) Predicted trajectories of tree community change in Amazonian rainforest fragments. Ecography, 40, 26–35.

Falkowski PG, Barber RT, Smetacek V (1998) Biogeochemical controls and feedbacks on ocean primary production. Science, 281, 200–206.

Field CB, Behrenfeld MJ, Randerson JT, Falkowski PG (1998) Primary production of the biosphere: Integrating terrestrial and oceanic components. Science, 281, 237–240.

Goldman JC, Glibert PM (1982) Comparative rapid ammonium uptake by four species of marine phytoplankton. Limnology and Oceanography, 27, 814–827.

Hemme D, Veyel D, Mühlhaus T et al. (2014) Systems-Wide Analysis of Acclimation Responses to Long-Term Heat Stress and Recovery in the Photosynthetic Model Organism Chlamydomonas reinhardtii. The Plant Cell, 26, 4270–4297.

Houlahan JE, McKinney ST, Anderson TM, McGill BJ (2017) The priority of prediction in ecological understanding. Oikos, 126, 1–7.

Hunter-Cevera KR, Neubert MG, Solow AR, Olson RJ, Shalapyonok A, Sosik HM (2014) Diel size distributions reveal seasonal growth dynamics of a coastal phytoplankter. PNAS, 111, 9852–7.

Hunter-Cevera KR, Neubert MG, Olson RJ, Solow AR, Shalapyonok A, Sosik HM (2016) Physiological and ecological drivers of early spring blooms of a coastal phytoplankter. Science, 354, 326–329.

Ishwaran H. and Kogalur U.B. (2017). Random Forests for Survival, Regression and Classification (RF-SRC), R package version 2.4.2.

Ishwaran H. and Kogalur U.B. (2007). Random survival forests for R. R News 7(2), 25–31.

Jochimsen MC, Kümmerlin R, Straile D (2013) Compensatory dynamics and the stability of phytoplankton biomass during four decades of eutrophication and oligotrophication. Ecology Letters, 16, 81–89.

Kehoe M, O’Brien K, Grinham A, Rissik D, Ahern KS, Maxwell P (2012) Random forest algorithm yields accurate quantitative prediction models of benthic light at intertidal sites affected by toxic Lyngbya majuscula blooms. Harmful Algae, 19, 46–52.

Kehoe MJ, Chun KP, Baulch HM (2015) Who Smells? Forecasting Taste and Odor in a Drinking Water Reservoir. Environmental Science and Technology, 49, 10984–10992.

Kremer CT, Williams AK, Finiguerra M et al. (2016) Realizing the potential of trait-based aquatic ecology: New tools and collaborative approaches. Limnology and Oceanography, 62, 253–271.

Litchman E, Klausmeier CA (2008) Trait-based community ecology of phytoplankton. Annual Review of Ecology, Evolution, and Systematics, 39, 615–639.

Magurran AE, Baillie SR, Buckland ST et al. (2010) Long-term datasets in biodiversity research and monitoring: assessing change in ecological communities through time. Trends in Ecology & Evolution, 25, 574–582.

McGill BJ, Enquist BJ, Weiher E, Westoby M (2006) Rebuilding community ecology from functional traits. Trends in Ecology & Evolution, 21, 178–85.

Mouquet N, Lagadeuc Y, Devictor V et al. (2015) Predictive ecology in a changing world. Journal of Applied Ecology, 52, 1293–1310.

Müller R, Pfeifroth U, Träger-Chatterjee C, Cremer R, Trentmann J, Hollmann R (2015) Surface Solar Radiation Data Set - Heliosat (SARAH) - Edition 1.

Paerl HW, Hall NS, Calandrino ES (2011) Controlling harmful cyanobacterial blooms in a world experiencing anthropogenic and climatic-induced change. Science of the Total Environment, 409, 1739–1745.

Paerl HW, Huisman J (2009) Climate change: A catalyst for global expansion of harmful cyanobacterial blooms. Environmental Microbiology Reports, 1, 27–37.

Petchey OL, Pontarp M, Massie TM et al. (2015) The ecological forecast horizon, and examples of its uses and determinants. Ecology Letters, 18, 597–611.

Pomati F, Jokela J, Simona M, Veronesi M, Ibelings BW (2011) An automated platform for phytoplankton ecology and aquatic ecosystem monitoring. Environmental Science Technology, 45, 9658–65.

Pomati F, Matthews B, Jokela J, Schildknecht A, Ibelings BW (2012) Effects of reoligotrophication and climate warming on plankton richness and community stability in a deep mesotrophic lake. Oikos, 121, 1317–1327.

Pomati F, Kraft NJB, Posch T, Eugster B, Jokela J, Ibelings BW (2013) Individual cell based traits obtained by scanning flow-cytometry show selection by biotic and abiotic environmental factors during a phytoplankton spring bloom. PLoS one, 8, e71677.

Rice EW, Baird RB, Eaton AD, Clesceri LS (eds.) (2012) Standard Methods for the Examination of Water and Wastewater, 22nd edn. American Water Works Association/American Public Works Association/Water Environment Federation.

Rivero-Calle S, Gnanadesikan A, Del Castillo CE, Balch WM, Guikema SD (2015) Multidecadal increase in North Atlantic coccolithophores and the potential role of rising CO2. Science, 350, 1533–1537.

Schulz J, Albert P, Behr H-D et al. (2008) Operational climate monitoring from space: the EUMETSAT satellite application facility on climate monitoring (CM-SAF). Atmospheric Chemistry and Physics Discussions, 8, 8517–8563.

Schwaderer AS, Yoshiyama K, de Tezanos Pinto P, Swenson NG, Klausmeier CA, Litchman E (2011) Eco-evolutionary differences in light utilization traits and distributions of freshwater phytoplankton. Limnology and Oceanography, 56, 589–598.

Smith VH (1983) Low nitrogen to phosphorus ratios favor dominance by blue-green algae in lake phytoplankton. Science, 221, 669–671.

R Core Team (2017). R: A language and environment for statistical computing. R Foundation for Statistical Computing, Vienna, Austria. https://www.R-project.org/.

Thomas MK, Kremer CT, Klausmeier CA, Litchman E (2012) A global pattern of thermal adaptation in marine phytoplankton. Science, 338, 1085–1088.

Thomas MK, Kremer CT, Litchman E (2016) Environment and evolutionary history determine the global biogeography of phytoplankton temperature traits. Global Ecology and Biogeography, 25, 75–86.

Thomas MK, Aranguren-Gassis M, Kremer CT, Gould MR, Anderson K, Klausmeier CA, Litchman E (2017) Temperature-nutrient interactions exacerbate sensitivity to warming in phytoplankton. Global Change Biology, 23, 3269–3280.

Williams JW, Jackson ST, Kutzbach JE (2007) Projected distributions of novel and disappearing climates by 2100 AD. PNAS, 104, 5738–42.

Zhu K, Chiariello NR, Tobeck T, Fukami T, Field CB (2016) Nonlinear, interacting responses to climate limit grassland production under global change. PNAS, 113, 10589–10594.

Zimmer A, Katzir I, Dekel E, Mayo AE, Alon U (2016) Prediction of multidimensional drug dose responses based on measurements of drug pairs. PNAS, 113, 10442–10447.

